# Altered dynamics may drift pathological fibrillization in membraneless organelles

**DOI:** 10.1101/598185

**Authors:** B. Tüű-Szabó, G. Hoffka, N. Duro, L. Koczy, M. Fuxreiter

## Abstract

Protein phase transition can generate non-membrane bound cellular compartments, which can convert from liquid-like to solid-like states. While the molecular driving forces of phase separation have been largely understood, much less is known about the mechanisms of material-state conversion. We apply a recently developed algorithm to describe the weak interaction network of multivalent motifs, and simulate the effect of pathological mutations. We demonstrate that linker dynamics is critical to the material-state of biomolecular condensates. We show that linker flexibility/mobility is a major regulator of the weak, heterogeneous meshwork of multivalent motifs, which promotes phase transition and maintains a liquid-like state. Decreasing linker dynamics increases the propensity of amyloid-like fragments via hampering the motif-exchange and reorganization of the weak interaction network. In contrast, increasing linker mobility may compensate rigidifying mutations, suggesting that the meshwork of weak, variable interactions may provide a rescue mechanism from aggregation. Motif affinity, on the other hand, has a moderate impact on fibrillization. Here we demonstrate that the fuzzy framework provides an efficient approach to handle the intricate organization of membraneless organelles, and could also be applicable to screen for pathological effects of mutations.

## Introduction

Protein self-assembly generates supramolecular complexes with a broad spectrum of structural and dynamical properties [1] (Figure 1). They range from amyloids, with solvent-excluded, solid cores, which are held together by interdigitating hydrogen bonds [2–4] to highly dynamical condensates [5, 6], with weak, interchanging, heterogeneous interactions [7, 8]. Signaling complexes can sample a wide regime between these states [9] to efficiently respond to noisy stimuli [10, 11] (Figure 1). Higher-order protein organization can generate stand-alone cellular species, which appear as puncta under the microscope [12]. These organelles, such as the nucleoli for ribosome biogenesis [13], Cajal bodies for spliceosome assembly, stress granules to store translationally arrested mRNA [14], P-bodies for mRNA turnover [5, 15], nuage granules in Drosophila germlines [16] all lack a membrane boundary.

**Figure 1.**
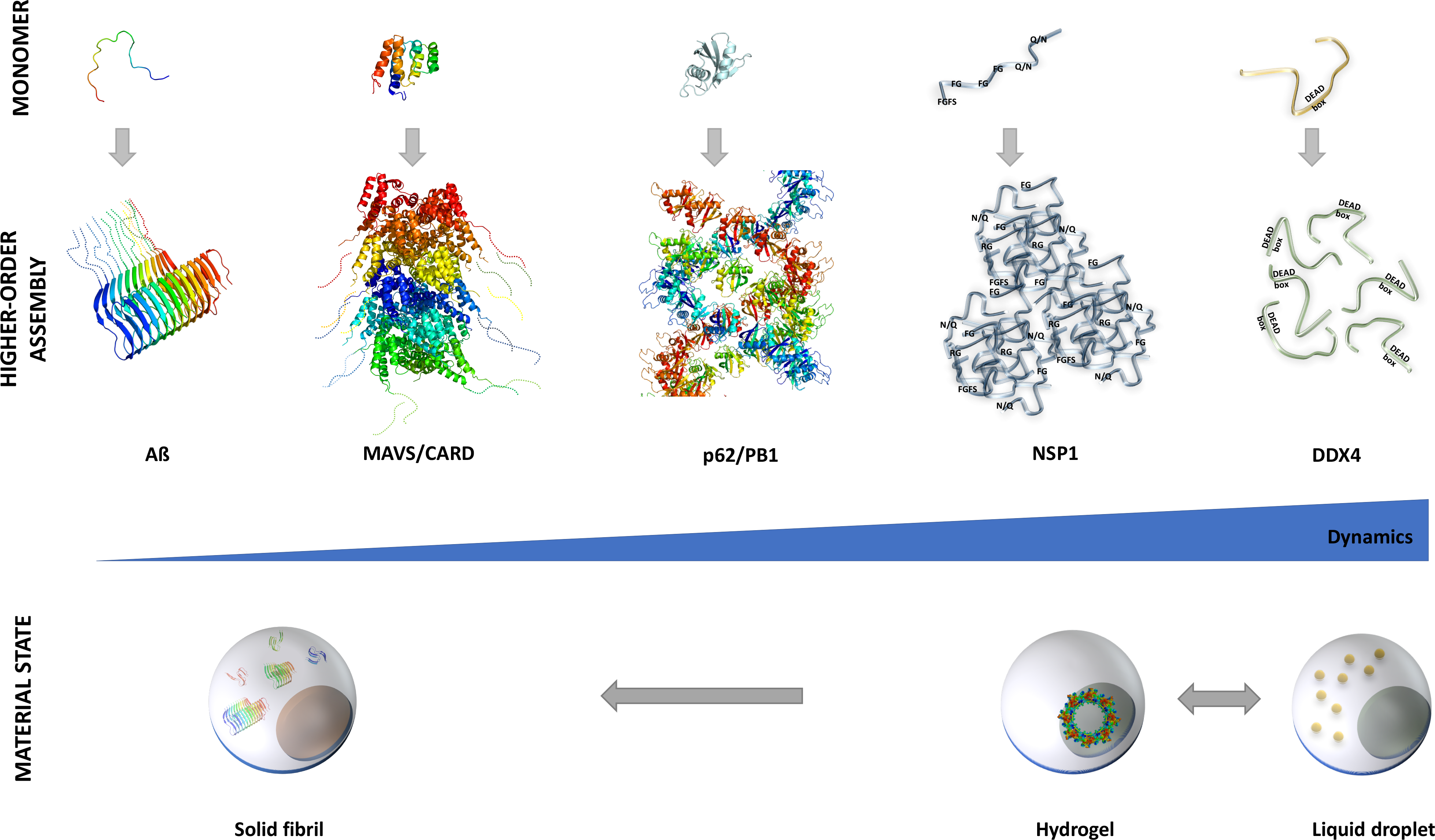
Structure and dynamics continuum of higher-order assemblies. The hierarchy of self-organization is shown from monomers to higher-order structures and cellular organelles (*from top to bottom*) Higher-order structures are displayed in the order of increasing dynamics and interaction heterogeneity *(from left to right)*: Amyloid β(PDB: 2mxu[78]), MAVS CARD domain (PDB: 2ms7 [79]), p62 PB1 domain (PDB: 4uf9 [80]), nucleoporin NSP1 [81], DDX4 nuage granule [16]. The conversion between liquid droplets and hydrogels is reversible, while transition to amyloids is an irreversible process. Formation of Intermediate states, like functional aggregates or dynamical signalosomes is also reversible.

Membraneless compartments function primarily by increasing concentration of the component proteins and the associated biomolecules (e.g. RNA) to impact biochemical reactions, to balance or store cellular factors, sequester toxic species, or increase avidity for low-affinity effectors [17]. Via all these means, protein-based organelles contribute to spatial and temporal regulation of a multitude of cellular processes, including protein transport [18,19], flux control [20] and responses to stress conditions [21]. Overall, membraneless compartments are highly dynamic cellular species to increase cellular fitness [22]. In accord, higher-order protein organizations often involve low-complexity (LC) and intrinsically disordered (ID) regions [23], which impart plasticity on the assembly.

Membraneless organelles are formed by phase separation [24] This process involves the exchange between intra- and intermolecular interactions of multivalent binding elements, which can be quantified by polymer theory [25, 26]. Membraneless organelles are held together by weak, multivalent contacts, where π-π and cation−π interactions play distinguished roles [8, 16, 27]. Multivalency is critical to generate reversible, cross-linked networks [28, 29], which might also involve weakly polar contacts [30]. For example, aromatic rings may interact in various orientations [31, 32], and can also act as H-bond acceptors [33], thus can simultaneously mediate contacts amongst multiple motifs. Linker regions connecting low-complexity motifs facilitate formation of many binding configurations via fast conformational exchange [7].

Membraneless organelles with highly dynamical or intrinsically disordered protein regions may exhibit liquid-like properties [34]. A series of evidence indicate that these protein-based organelles are metastable and tend to convert towards more solid aggregates [35–37]. Formation of amyloid-like fibrils is often associated with a variety of pathologies [38], such as amyotrophic lateral sclerosis (ALS), frontotemporal dementia (FTD), inclusion body myopathy (IBM) [39]. Inherited forms of these degenerative diseases are caused by missense mutations in proteins such as hnRNPs [40], Fus [37], TIA-1 [41] or TDP-43 [42]. Posttranslational modifications may also contribute to aggregation or malfunction [43]. Biophysical studies indicate that single-point mutations, such as D262V in hnRNPA1 [36] or G156E in Fus [37] induce a gradual transition of the liquid droplet towards solid fibrils, similarly to the effect of aging [17]. The molecular driving forces leading to aggregation however, are remained to be elucidated.

Here we aim to describe the material-state conversion of membraneless organelles from highly dynamic to solid-like states using a recently developed simulation methodology. The approach accounts for multiple, weak contacts between tandem motifs, and it has been applied to describe phase transition of generic models [44]. In this study, we introduced mutations into the droplets and followed the trajectories of fibril-formation. These mutations systematically affected linker dynamics and motif affinity. First, we related phase transition to linker flexibility/mobility, and showed that cooperative network of weak contacts promotes self-organization into large polymers. Then we related the propensity of amyloid-like elements and entropy of the system to motif-affinity and linker dynamics. We found that more rigid linkers promote fibrillization by slower exchange between multivalent motifs. Furthermore, increasing linker dynamics may also compensate the effect of rigidifying mutations. Interestingly, changes in motif affinity have moderate impact on aggregation. Therefore, we propose that the meshwork of weak, variable interactions may compensate pathological mutations and prohibit conversion to solid-like fibrils.

## Methods

### Model system

#### Model protein

The model comprised tandem repeats of interacting motifs (7 AA) and linkers (10 AA) (Figure 2A). A 75 AA model protein with 5 binding elements or motifs (5 × 7 AA), and 4 linkers (4 × 10 AA) has been selected based on previous experimental [29] and computational results [44]. Each amino acid was characterized by two values: binding affinity (s) and dynamics (d). Binding affinity was related to the strength/weakness of motif interactions, defined on a relative [0,1] scale. This value reflects the binding preference of a residue (i.e. likelihood to be engaged in interactions), which could be estimated from dissociation constants (K_D_) or computed binding free energies (ΔG_bind_).

**Figure 2.**
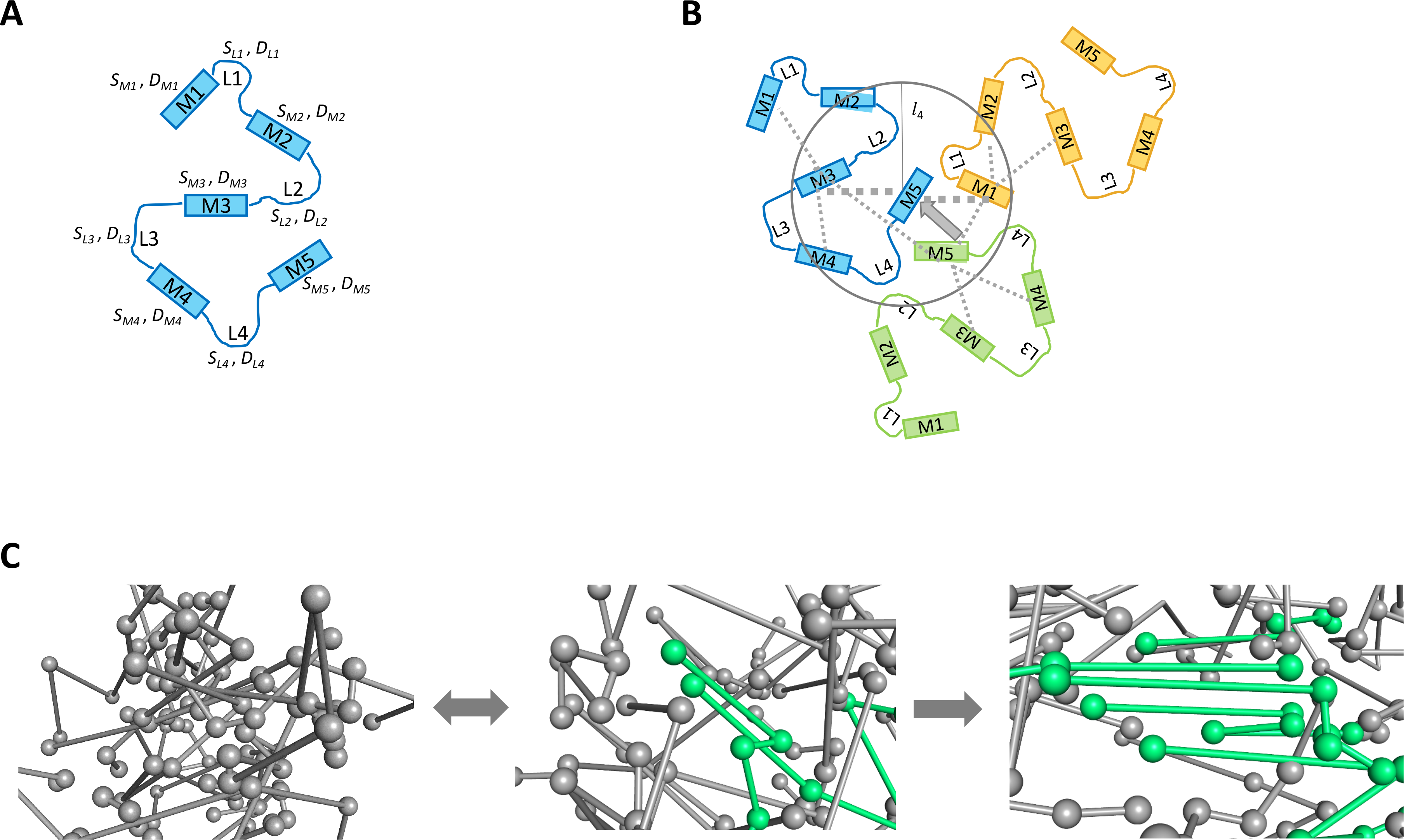
Model system. **(A)** Generic, multivalent model with 5 binding motifs (M1-M5), which are connected by 4 linkers (L1-L4). Each element is characterized by interaction affinity (S_i_) and dynamics (D_i_). **(B)** Multisite interactions in the multivalent model. Here only 3 molecules are shown. Binding motifs within the gray cirle influence M5 (blue) interactions. The radius is set according to the length of L4 (*l_4_*). The widths of the dashed lines indicate different coordination spheres. **(C)** Reorganization of heterogeneous polymers (gray) to amyloid-like elements (lime) (*from left to right*). Configurations are derived from the simulations, spheres represent the motifs, and sticks correspond to linkers. Linkers are not straight in the simulations, they can adopt numerous topologies and conformations based on their D (dynamics) value.

Affinity of an interaction element/motif (α_i_) was obtained as an average of the residue-based values:

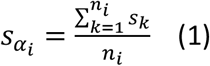

where *s*_*k*_ is the binding affinity of residue *k* and *n*_*i*_ are the number of residues in the interacting motif.

Dynamics referred to the rate of conformational exchange in the assembly. This is often difficult to quantify experimentally [7], especially in case of transient interactions [45, 46]. Thus, we used a simplified [0,1] scale for characterizing dynamics, reflecting to what extent the mobility of the given residue is retained in the bound-state. For example, D = 0 corresponds to a fully structured residue with a well-defined interaction in the assembly, and D=1 to a highly dynamic residue with constant exchange between different binding modes in the presence of the partner [47]. Those residues, which may fold upon binding under a specific context, usually sample the D=0.3-0.7 regime. Dynamics could be estimated using NMR datasets [48] or via bioinformatics predictions [49].

Linker dynamics was defined as the average of the residue-based values:

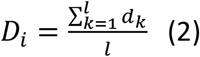

where *d*_*k*_ is the dynamics of residue *k*, and *l*_*i*_ is the length of the linker.

The initial values for binding affinity were set to 0.5 for residues in the interacting motifs and 0.05 for residues in the linkers. Dynamics was set to 0.05 for residues in the motifs and 0.3 for residues within the linkers. Both parameters were systematically varied by mutation effects. Rigidifying mutations decreased dynamics of linker L3, between M3 and M4 motifs from 0.3 to 0.05 in 0.05 steps (Figure 2A). To probe the impact of linker topology, a similar procedure was carried out for L1 between M1 and M2 motifs. Mutations affecting motifs decreased binding affinity of M3 from 0.5 to 0.2 in 0.1 increments (Figure 2A, 2B).

#### Simulated system

The system consisted of 1000 identical molecules (75 AA each), which were placed randomly into 343 cubic boxes with a length of 50 residues. No steric overlap was allowed. In this coarse-grained representation, water was not explicitly considered.

## Definitions

### Binding of two molecules

Two molecules were associated if at least 1 interaction was established between two motifs, which were located on separate molecules.

### Oligomers

Oligomers contained (1 < n < 25) molecules, each of which was connected to the rest of the cluster via at least 1 motif-interaction.

### Polymers

Polymers contained n ≥ 25 molecules, each of which was connected to the rest of the cluster via at least 1 motif-interaction.

### Aggregates/fibrils

Amyloid-like elements were defined based on a regular interaction pattern amongst n ≥ 5 molecules. Regular interaction patterns were connections between at least 2 consecutive binding elements on both interacting partners (Figure 2C).

## Theoretical framework

### Interactions between two motifs

Binding affinity between two motifs was characterized by the average of the individual binding preferences:

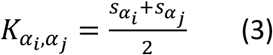

where 𝑠_*α*_*i*__ and 𝑠_*α*_*j*__ are the binding preferences of the interacting α_i_ and α_j_ motifs. Two motifs were bound if *K*_*α*_*i*_,*α*_*j*__ > 0.2.

### Potential interactions with multivalent sites

In the fuzzy framework, partial interactions with multiple sites are also considered. First, we define the volume, within which potential binding elements were located (Figure 2B):

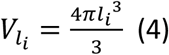

where l_i_ is the length of the linker, and the spherical volume *V*_*l*_*i*__ is centered on the binding element α_i_.

Then the number of binding elements, which were located within this volume was determined (Figure 2B):

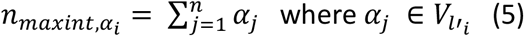

where *n*_*maxint,α*_*i*__ is the number of binding motifs, which have at least 1 residue within the defined volume. *n*_*maxint,α*_*i*__ is the maximum number of the potential interaction sites, which are available for interactions with α_i_ motif. We reasoned that the number potential binding sites also modulate the interaction between two motifs (α_i_, α_j_).

### First association event (anchoring step)

The association probability in the first step was defined as:

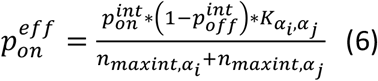

where *K*_*α*_*i*__,_*α*_*j*__ is the binding affinity between α_i_ and α_j_ motifs (eq. 3), *n*_*maxint,α*_*i*__ and *n*_*maxint,α*_*j*__ are the maximum number of potential interaction elements around α_i_ and α_j_ motif, respectively. Qualitatively, competing motifs reduce the association probability between two motifs (α_i_, α_j_) (Figure 2B). 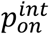 and 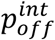 is the intrinsic association and dissociation probability, which were set according to the reference [29]. The binding was realized if 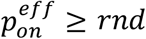 where *rnd* was a random number.

### Determination of occupancy

After the first binding step, an occupancy of each binding element was determined. As the applied framework also considers partial interactions, a fuzzy union (s-norm) operator (based on a fuzzy set theory [50]) for each connected motif had to be determined:

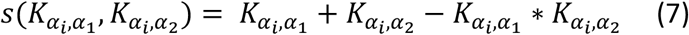

where *K*_*α*_*i*__*K*_*α*_*i*__ is the binding affinity of α_i_ and α_j_ motifs (eq. 3). The occupancy of each motif was defined as an algebraic sum:

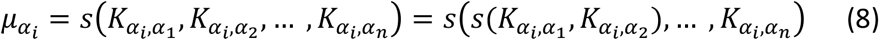

where *s*(*K*_*α*_*i*_,*α*_*1*__, *K*_*α*_*i*_,*α*_*2*__, … *K*_*α*_*i*_,*α*_*n*__) is the fuzzy union operator in eq. 7. Occupancy (eq. 8) is computed for each, even partially bound motifs (*K*_*α*_*i*_,*α*_*1*__ > 0.2). This formalism has been developed to describe simultaneous, weak interactions in a multivalent system (eg. cation-π and π - π involving the same tyrosine residue).

### Subsequent binding events

#### Modified binding affinity

Once two motifs were attached, the binding affinity between two motifs (eq. 3) was modified by the corresponding occupancies of potential binding sites (Figure 2C):

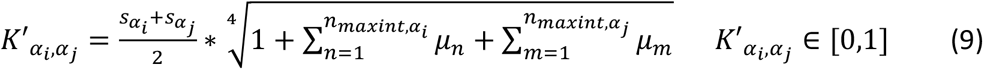

where *μ*_*n*_ and *μ*_*m*_ are the occupancies of all potential binding elements (eq. 5) within the given volume (*V*_*l*_*i*__) around α_i_ and α_j_ (eq. 4). Thus, high occupancies of the potential interaction motifs within *V*_*l*_*i*__reduce the competition for α_i_ and α_j_, respectively, and thus increase *K*_*α*_*i*__, _*α*_*j*__affinity.

#### Modified association probability

After the first binding event (once the two motifs were bound) the association probability was also modified by the corresponding occupancies:

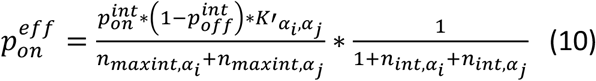

where *K′*_*α*_*i*__, _*α*_*j*__ is the modified binding affinity defined in eq. 9, *n*_*int,α*_*i*__ and _*int,α*_*j*__ are the actual number of binding elements, which are bound to α_i_ and α_j_, respectively.

#### Dissociation

The dissociation probability had an inverse relationship to the binding affinity:

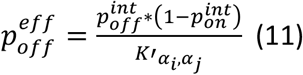

where the intrinsic association 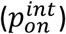 and dissociation 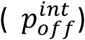 probabilities are the same as in eq. 6, and *K′*_*α*_*i*__, _*α*_*j*__ is the modified binding affinity defined in eq. 9.

#### Diffusion

During the simulation, the different kinds of species (individual molecules, oligomers and polymers) could be transformed into one of the neighboring boxes with a probability:

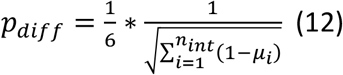

where 1 − *μ*_*i*_, is the available interaction capacity of a given α_i_ binding element. Summation is carried out for all motifs, which are bound to α_i_ (*n*_*int,α*_*i*__). Therefore, larger polymers with more available binding capacity are less likely to move to another box. During diffusion, the molecules/oligomers/polymers were moved as rigid units.

#### Accounting for linker dynamics in fuzzy models

The advantage of using a fuzzy model, that it can describe multivalent/multisite binding. The number of potential interaction sites (eq. 5) were determined within the *V*_*l*_*i*__ volume (eq. 4), which was defined by the linker length as a radius. In the fuzzy model, the association probability (eq. 10) and binding affinity (eq. 9) between two motifs were dependent on *n*_*maxint,α*_*i*__(eq. 5) as well as on the occupancies of the potential binding sites (eq. 8), so they are indirectly influenced by the linker properties. If the linker is infinitively flexible (D_i_=1, eq. 2), the neighboring motifs can sample the whole volume (*V*_*l*_*i*__, eq. 4), which is available for interactions (Figure 2B). Less dynamical linkers, however, cannot visit the whole space for interactions, as the binding elements become restricted. Thus, if *D*_*i*_ < 1, the linker length has to be modified to take the reduced dynamics into account:

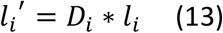

where *D*_*i*_ is the linker dynamics and *l*_*i*_ is the length of the linker. *l*’_i_ is used to obtain the volume available for interactions (eq. 4).

### Computational protocol

In the first simulation step, binding between multivalent motifs could take place according to the association probabilities (eq. 6). Once the initial contacts were made, the occupancies were determined in each step. The binding affinities (eq. 9) and association probabilities (eq. 10) were modified accordingly. In all subsequent steps, binding or dissociation of an interaction element from the other available or bound motifs were decided based on the association (eq. 10) and dissociation probabilities (eq. 11). Each species: individual molecules, oligomers and polymers were moved to the one of the neighboring boxes according to the diffusion probabilities (eq. 12). The simulation steps are given in arbitrary units. Using a contour length of 4 Å/residue [51], and an approximate diffusion constant D ~ 1.1 × 10^−6^ cm^2^s^−1^ [52], a simulation step approx. corresponds to a μs motion (rough estimate for individual molecules).

The simulation consisted of two regimes. In the first period (step = 0 – 10000) oligomers and large polymers formed and their propensity converged within ± 2.5 % (phase transition). In the second period (step = 10000 – 15000) large polymers were in equilibrium with individual molecules and oligomers, but the interaction patterns started to reorganize (Figure 2C). Such reorganization also involved a slow conversion towards more stable, regular patterns, referred to as aging or fibrillization. Single point mutations affecting dynamics or affinity were introduced into L3 (between M3 and M4 motifs) or L1 (between M1 and M2 motifs) linkers, or within M3 motif (Figure 2A, 2B) in the beginning of the second regime (step = 10000). Linker mutations decreased linker mobility, whereas motif mutations decreased binding affinity as described above. Simulations were conducted until step = 15000 for both native and mutant sequences and the system composition was analyzed in this part of the trajectory (step = 10000 – 15000).

### Analysis

In each simulation step, the composition of the system was determined according to the definitions as given above. The fraction of oligomers and polymers versus individual molecules were computed. Polymers were further analyzed for amyloid-like elements; interactions between consecutive binding elements involving n ≥ 5 molecules within the polymer.

The entropy of the system was computed based on Shannon-entropy, which was modified for the fuzzy framework [53]:

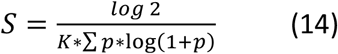

where K=1/n, and *n* is the number of the polymers with different topologies); and *p* is the propensity of the polymer with a given topology. The entropy formula was scaled to [0,1].

## Results

### Phase transition in fuzzy and non-fuzzy models

As the application of fuzzy models is relatively new to biology [54, 55], we highlight a few distinctions between the fuzzy and non-fuzzy models. In non-fuzzy models the interacting motifs are fully engaged in binding with a unique target, *K*_*α*_*i*__, _*α*_*j*__ = 1 (eq. 3), while in fuzzy models a partial binding (*K′*_*α*_*i*__, _*α*_*j*__ < 1, eq. 9) to multiple sites is also considered [44]. This corresponds to multiple, weak contacts (e.g. π − π and cation-π, which may also take place simultaneously (Figure 2C). Weak interactions however, are less likely to occur as the association probabilities (eq. 6, 10) are proportional to binding affinities (*K*_*α*_*i*__, _*α*_*j*__ eq. 3 and *K*_*α*_*i*__, _*α*_*j*__ eq. 9). The dissociation probabilities (eq. 11) are also higher for weak interactions, so partial contacts in the fuzzy model are more likely to dissociate than the full binding in the non-fuzzy model. Therefore, simulations using fuzzy models could only produce large polymers (n ≥ 25) in case of multivalent systems with n ≥ 4 [44].

Phase transition is highly sensitive to protein dynamics and increased linker mobility facilitates the process [34, 56, 57]. In accord, models with linker dynamics of D=0.40 and 0.50 were almost fully polymerized; the propensity of polymers (n ≥ 25) exceeded 90 % (step = 9000 – 10000 steps, Figure 3A, 3B). Fuzzy and non-fuzzy simulations have converged to similar results, but the rate of polymerization was approximately two times faster using the fuzzy approach (Figure 3A, 3B). Models with lower dynamics (D = 0.20) markedly deviated in the fuzzy and non-fuzzy simulations: while 62 % of the molecules belonged to polymers in the fuzzy simulation, only 16 % of the molecules self-assembled in the non-fuzzy treatment (Figure 3A, 3B). These observations underscore the role of linker dynamics in facilitating phase transition via organizing the network of weak, possibly partial contacts in multivalent systems.

**Figure 3.**
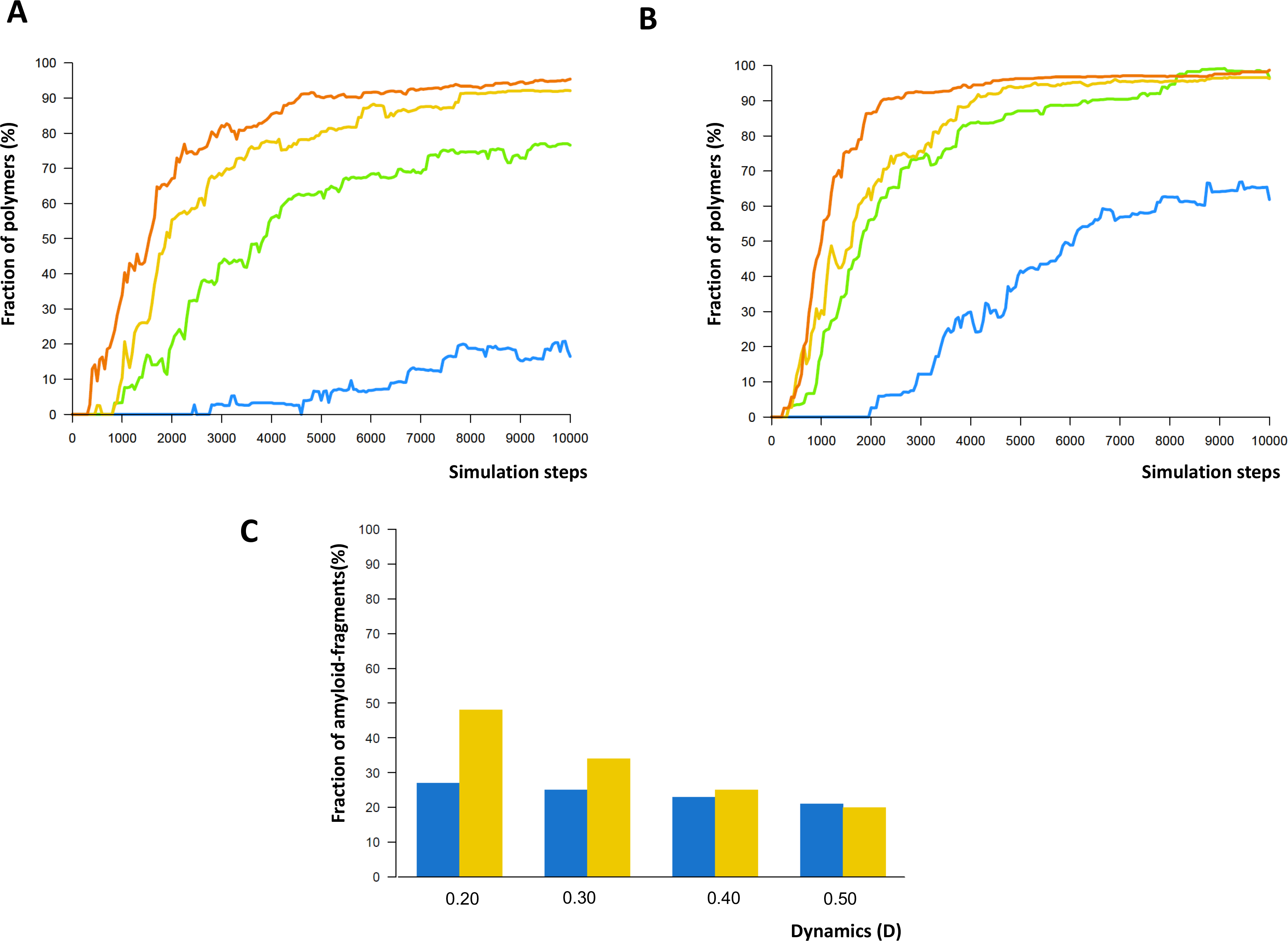
Phase transition in (A) non-fuzzy and (B)fuzzy models. Sequences with different linker dynamics are displayed by orange (D=0.50), yellow (D=0.40), lime (D=0.30), marine (D=0.20). **(C) Amyloid-like fragments in polymers.** Non-fuzzy (blue) and fuzzy models (gold) models with different linker dynamics are shown.

### Reorganization of higher-order assemblies in fuzzy and non-fuzzy models

Self-assembly of molecules into large polymers generates highly heterogeneous interaction patterns, which dynamically rearrange according to the association and dissociation probabilities (eq. 10, eq. 11). During this process, the fuzzy approach takes the effect of multivalent motifs into account via considering weak, partial interactions with multiple potential binding sites. Association probability between two motifs (eq. 10) is decreased by already bound partners as well as potentially competing sites; while increasing occupancy (eq. 8) of competing sites improves affinity (eq. 9). Dissociation between two motifs (eq. 11) is enhanced by already bound partners and competing motifs with low occupancy (eq. 8). The polymers can also exchange molecules according to the diffusion probabilities, which is disfavoured by higher interaction capacity (eq. 12).

During the reorganization process, heterogeneous interaction networks can also rearrange into more ordered patterns, where consecutive binding elements establish contacts with each other (Figure 2C). Such zipper interactions may also be present in membraneless organelles [58]. In accord, 21 – 27 % of polymers contained amyloid-like elements (n ≥ 5 molecules) in the simulations with the non-fuzzy model (Figure 3C). In the fuzzy simulations, the propensity of zipper-containing polymers varied between 21 – 47 % depending on linker dynamics. At lower linker dynamics (D=0.20) considerably higher propensity of regular interaction patterns (i.e. amyloid-like elements) were observed in the fuzzy than in the non-fuzzy approach, whereas with more mobile linkers (D=0.50) the results of the two simulations were comparable (Figure 3C). Thus, the fuzzy model appears to be more sensitive to linker dynamics, which is a critical factor in driving the rearrangement of motif-interactions.

### Linker mutations may drift fibrillization of membraneless organelles

Single-point mutations may accelerate conversion of liquid-like membraneless organelles to solid fibrils [36, 37, 39]. After the propensity of polymers had converged in our simulations (step = 10000), the dynamics of the linkers and the binding affinity of motifs were affected by mutations and the impact on polymer organization was studied. The dynamics of L3 (Figure 2) was systematically reduced from 0.30 to 0.05 in 0.05 steps, while the mobility of the other linkers was kept constant (D=0.30). A reduction in L3 linker dynamics from D=0.30 to D=0.20 increased the propensity of polymers with amyloid-like elements from 34.0 % to 46.0 % in fuzzy simulations (steps = 10000 – 15000), as compared to the total number of polymers (Figure 4A). A further decrease in L3 dynamics to D=0.05 raised the fraction of amyloid-containing polymers to 56.0 % (Figure 4A). Linker mutations in L1 (between M1 and M2 motifs) were much less effective in inducing fibrillization, as only 42 % of the polymers contained amyloid-like elements (Figure 4B).

**Figure 4.**
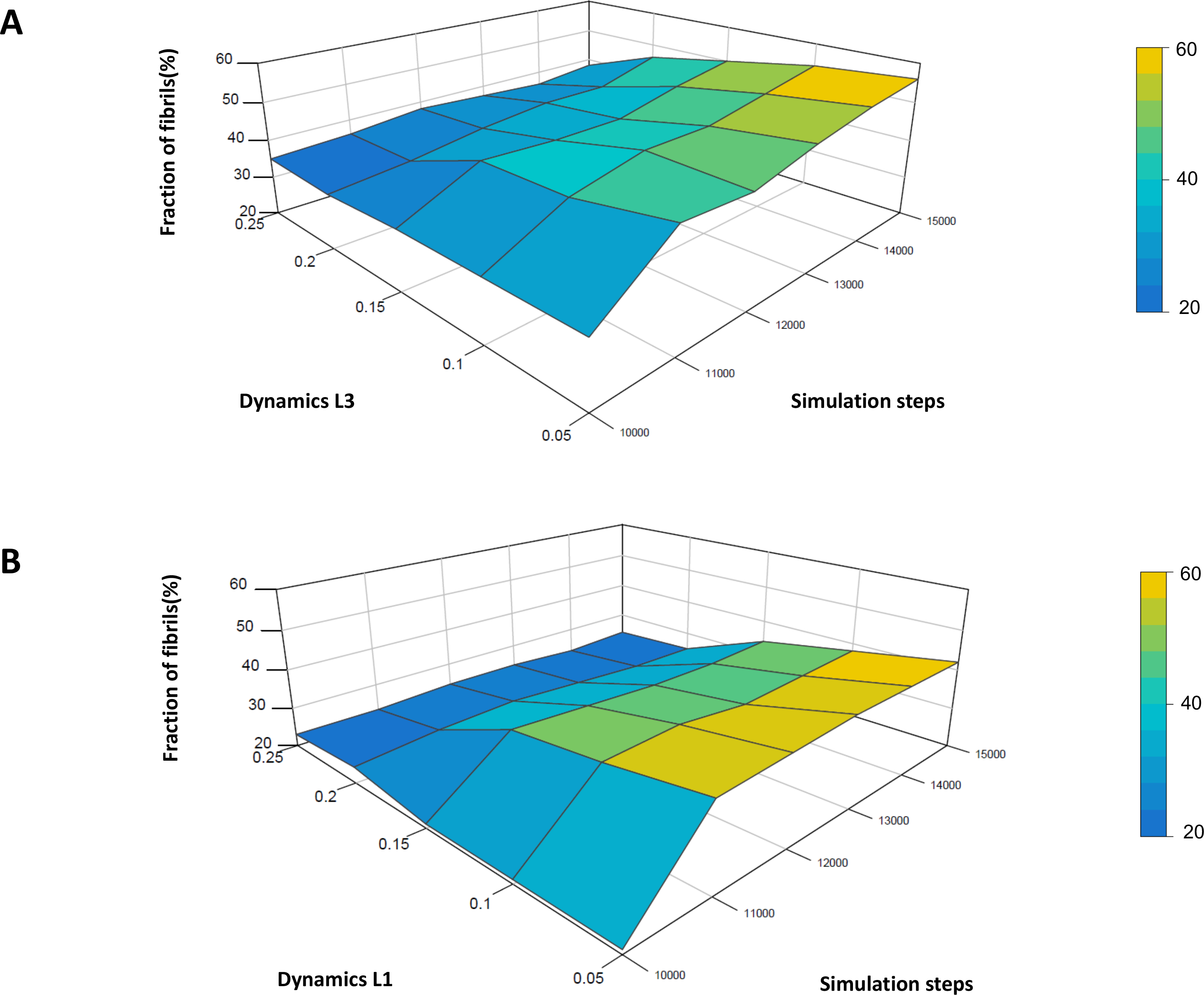
Fibrillization induced by rigidifying the (A) L3 and (B) L1 linker. The propensity of polymers with amyloid-like elements is shown as a function of affected linker dynamics during step= 10000 – 15000 of the trajectory. Dynamics of all other linkers was set to D=0.30.

Fibril formation decreases the entropy of the system leading to more solid-like states [41]. A modified Shannon-entropy [53] (eq. 14) was used to compute entropy in the fuzzy simulations. Decreasing L3 dynamics from D=0.30 to D=0.20 reduced entropy by 34.8 % as compared to its value in the native state (D=0.30 for all linkers, step = 10000). Further decrease in linker mobility (to D=0.05) resulted in an entropy-drop by 69.6 % as compared to the native state (Figure 5A). In case of L1 mutations (D=0.05), entropy was reduced only by 28.3 % as compared to the native state (Figure 5B). In contrast to linker dynamics, mutations decreasing binding site affinity had a negligible impact on the fraction of fibrils and system entropy (Figure S1, Figure 6B). These observations demonstrate that linker dynamics is a major factor in regulating the exchange of the multivalent binding motifs, slowing down of which leads to more ordered interaction patterns and fibril propagation.

**Figure 5.**
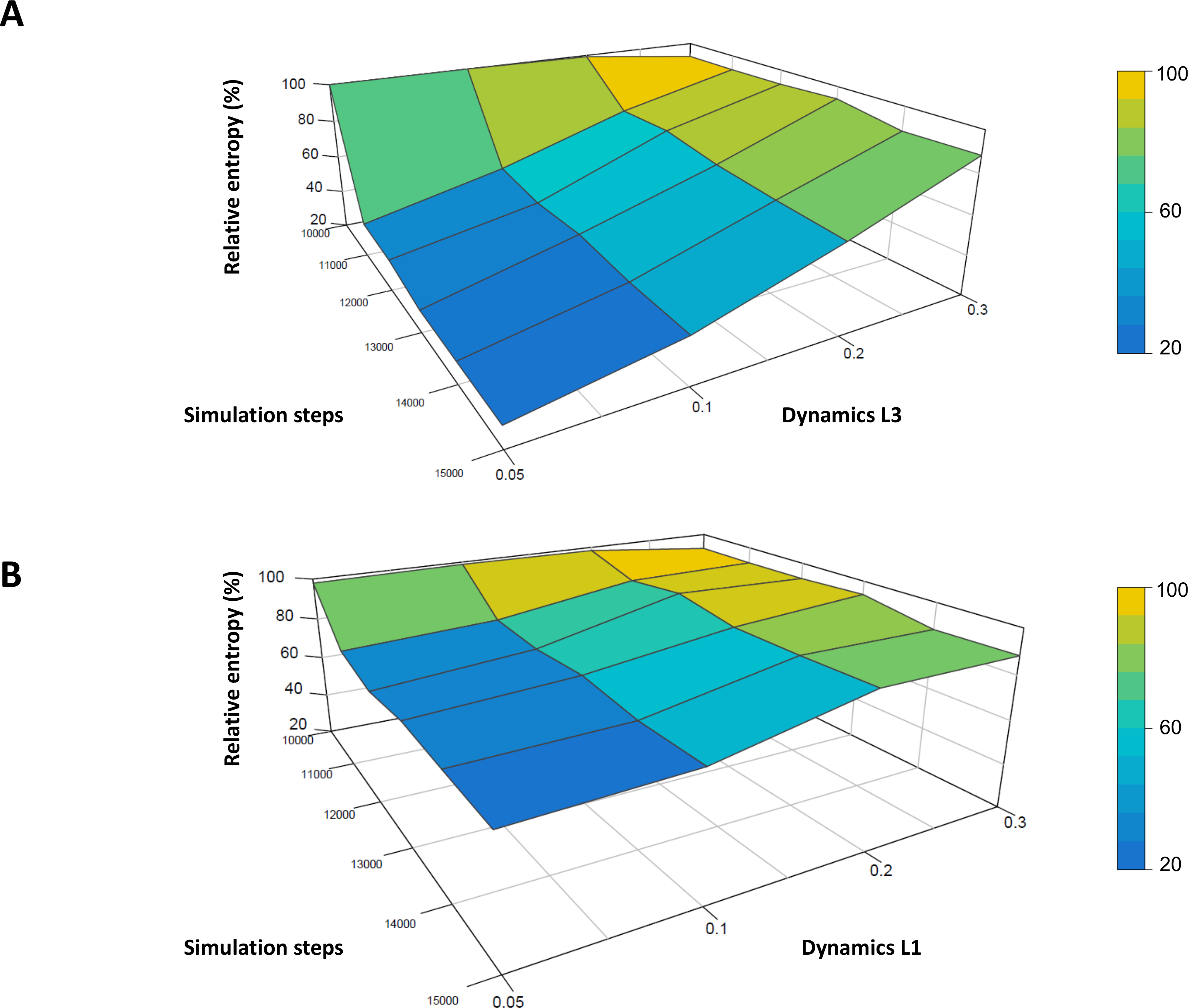
Relative entropy upon rigidifying the (A) L3 and (B) L1 linker. The entropy relative to the native state (step = 10000) is shown as a function of affected linker dynamics during step= 10000 – 15000 of the trajectory. Dynamics of L3 was decreased to D=0.10 at step = 10000 of the simulation, while D=0.30 was kept for all other linkers.

**Figure 6.**
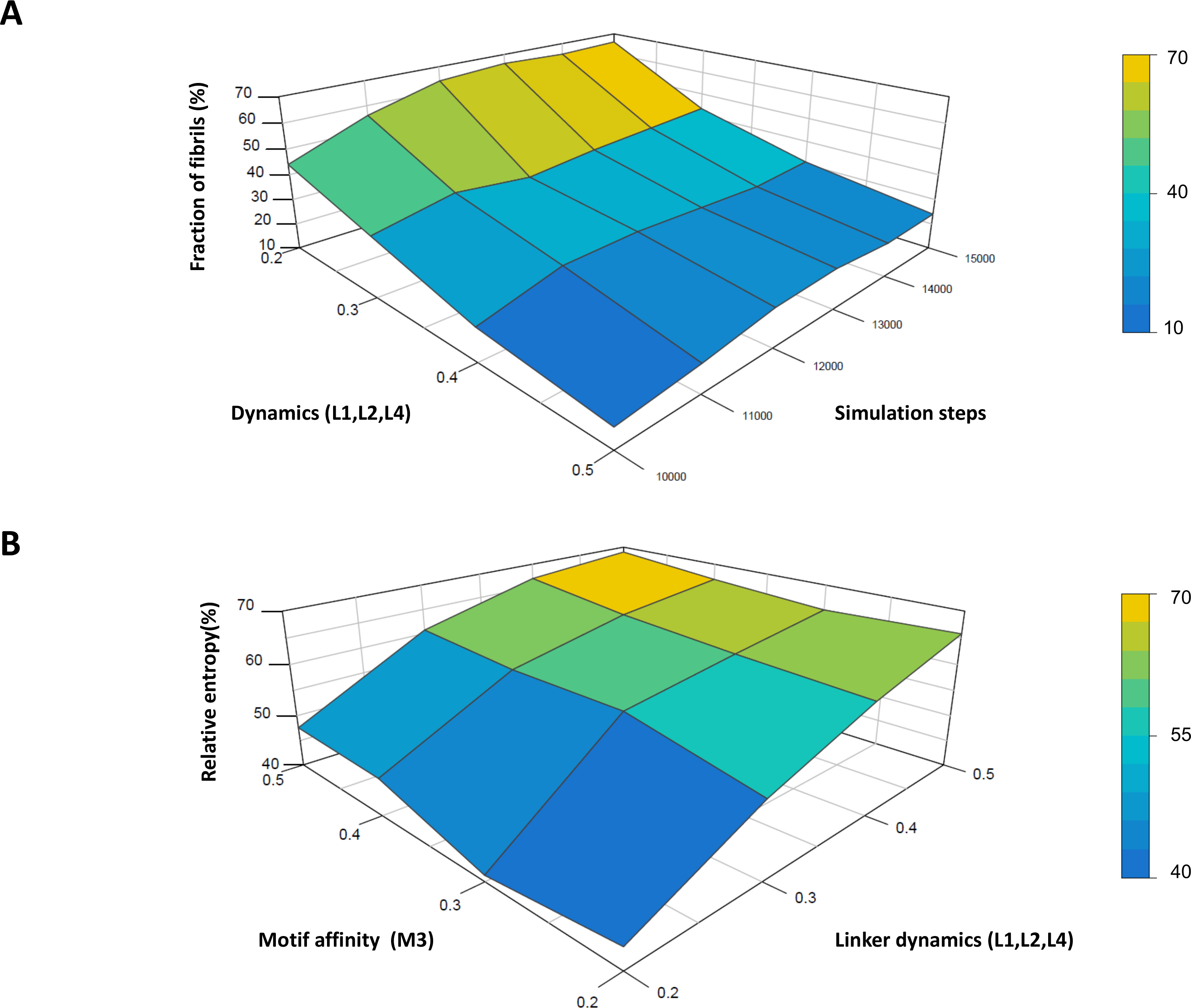
Increased dynamics may block fibrillization. **(A)** The propensity of polymers with amyloid-like elements is shown as a function of dynamics during step= 10000 – 15000 of the trajectory. Dynamics of L3 was decreased to D=0.10 at step = 10000 of the simulation, while dynamics of the other linkers was kept constant. **(B)** The entropy (step=15000) relative to the native state (step = 10000) is shown as a function of L1, L2, L4 linker dynamics and M3 motif-affinity. Dynamics of L3 was decreased to D=0.10 at step = 10000 of the simulation, while dynamics of the other linkers was kept constant. Affinity of all other motifs was set to S=0.50.

### Cooperativity and increased dynamics may provide a compensatory mechanism from aggregation

The distinct behaviour of L1 and L3 mutations indicates an interplay between the multivalent elements during the reorganization of polymers and material-state conversion. This raises the question on possible compensatory mechanisms via cooperative effects. To probe this scenario, linker flexibilities were systematically varied between D=0.20 to D=0.50, while rigidifying mutations were introduced into the L3 linker (D=0.1, step = 10000). Stiff linkers (D=0.20) were prone to aggregate even without mutations, which further increased the propensity of amyloid-like elements by 22 % (steps = 10000 – 15000, Figure 6A). In contrast, increasing flexibility of the non-mutated linkers (L1, L2, L4) considerably reduced the fraction of fibrils. Rigidifying L3 mutations lead to only 7 % enrichment in amyloid-like elements using models with D=0.50 (L1, L2, L4). Reducing M3 motif affinity had smaller impact on fibril formation than modulating linker dynamics Figure 6B). These results indicate that the network of weak, partial contacts may compensate the effect of rigidifying mutations and impede aggregation (Figure 7).

## Discussion

Material state conversion is a central problem in membraneless organelles [59]. Experimental data suggests that dynamic, weak, motif-based multivalent interactions are intrinsically metastable until fully transformed to the amyloid state [60, 61]. Crystal nucleation driven by short-range attractive forces [35] is analogous to this process, which may be also preceded by liquid-liquid phase transition. It has been a long-standing question to what extent the molecular structure in liquid droplets resembles to those in solid fibrils? Independent lines of experimental evidence indicate that identical interactions underlie both types of assemblies [60, 62, 63]; amongst which πcontacts are distinguished [8]. Low-complexity aromatic-rich kinked segments (LARKs, [64]) may be represented as labile β-elements in both liquid and solid states [58], which could be cross-linked similarly to hydrogels [65]. Liquid droplets exhibit multiple, alternative cross-linking arrangements, which may vary according the context or post-translational modifications [21, 66, 67]. In contrast, fibrils have reduced number of crosslinking microstates leading to a significant decrease in entropy [4, 37].

The lifetimes of these contacts however, may significantly differ between different assemblies with distinct physical states [1]. While interactions are transient and highly heterogeneous in liquid droplets [7], they are more stable and permanent in hydrogels and fibrils [62, 65]. The dynamics of the interactions could be qualitatively characterized by FRAP studies, which estimates reorganization of ensembles of interacting motifs, rather than quantify the lifetimes of the individual contacts. The latter could be derived from NMR studies, mostly R_2_ relaxation rates, which indicate μs-ms motions for dynamical interaction profiles [68]. In liquid droplets local motions also appear to be preserved in the bound form [7]. Linker dynamics should also be related to other physical properties, such viscoelasticity of the droplets, which was not addressed in this study.

Here we used a novel simulation approach to investigate the effect of linker dynamics in phase transition and material state conversion. The applied model aimed to recapitulate various aspects of multivalent systems. Most importantly, it assumed that weak, polar interactions, such as π − π or cation-π contacts may occur simultaneously forming a labile network in the multivalent system. These simultaneous interactions were represented by partial contacts, which mutually influenced binding characteristics. Weak contacts were more difficult to form and easier to separate, similarly to transient interactions. In this formalism, occupancies of the multivalent binding sites contributed to modulating associations/dissociations of two motifs.

In addition, we demonstrated that the network of weak contacts could facilitate phase transition, especially in case of more flexible/dynamical linkers. We reason, that continuous reorganization of such weak interactions increases the number of microstates and imparts fluidity on the liquid droplets. Along these lines, local decrease in linker dynamics may reduce the rate of exchange and bias for more ordered interaction patterns. Indeed, mutations rigidifying L3 significantly increased the propensity of polymers with amyloid-like elements. In accord to experimental data, droplets in our simulations contained a small fraction of fibril-like fragments (i.e. molecules connected via consecutive motifs). In case of flexible or disordered linkers (D ≥ 0.30) unidirectional propagation leading to fibrils did not occur, as the interaction topologies between these amyloid-like elements could be reorganized. This again points to the importance of linker dynamics in determining the material-state of droplets or condensates. Along these lines, mutations reducing linker dynamics bias towards more stable, regular interactions and drift the system towards aggregation. Concomitantly, entropy decreases and solid-states are likely to be formed.

These results are in line with earlier observations on amyloid β(Aβ) aggregation. Molecular dynamical studies showed that antiparallel Aβoligomers are mobile and undergo significant conformational rearrangements to accommodate the monomers [69]. This mechanism, referred to as ‘dock– lock’, comprises two steps: first docking the unfolded monomer to the preformed fibrils, and then structural conversion of the oligomers. Although the antiparallel cross-βstructure is stable, the oligomers also visit alternative structures and exhibit extensive fluctuations on hundred ns timescale. Antiparallel order increases with peptide size, which accelerates propagation of the fibrillar structure [69].

Different linkers cooperate in regulating the reorganization of weak, multivalent motifs. This can be illustrated by the distinct effects of rigidifying L3 or L1 linkers, respectively. Stiffening L3 was more effective in promoting fibrillization than those in L1 linker. Increased linker dynamics in non-mutated linkers also regulate reorganization of weak motifs, thereby offer a compensatory mechanism to rigidifying mutations. Increasing linker flexibility from D=0.20 to D=0.50 significantly decreased the propensity of polymers with amyloid-like elements, which were induced by L3 linker mutations (steps = 10000 – 15000). These results suggest that reorganization of weak, polar contacts may provide a rescue mechanism from aggregation.

Ultimately, how realistic is this model with a simplistic representation of multivalent molecules (5 motifs and 4 linkers)? Artificially engineered multivalent systems have been successfully used as models of phase transition [29, 56, 57]. The critical element of the applied framework is to consider motif-interactions with multiple sites. Experimental and computational studies highlight that ID binding is largely influenced by non-native interactions [70–72]. Even in case of templated folding, non-native contacts frustrate the binding free energy landscape [73, 74], and enable multiple bound configurations [75]. These effects have been largely overlooked thus far in interactions of flexible or ID proteins [47]. A series of experimental evidence, however, support the functional importance of the network of weak polar contacts in highly dynamic systems [76, 77]. The mathematical framework, based on the fuzzy set theory is applicable to capture these cooperative effects [55], which can importantly contribute to both phase transition and fibrillization.

## Conclusion

A recently developed computational approach has been applied to simulate how pathological mutations induce fibrillization in membraneless organelles. This framework enabled to study how the weak, interaction network of multivalent motifs influence the structural and dynamical properties of the system. We demonstrate that linker flexibility has a critical role in determining the material-state of membraneless organelles: increased dynamics facilitates phase transition, while it impedes the conversion to aggregates. Linker dynamics is a major factor in reorganizing the weak interaction meshwork. We show that simultaneous, partial contacts (e.g. π-πand cation-π interactions via the same tyrosine residue) increase heterogeneity and contribute to reducing fibrillization. Rigidifying linker mutations slow down the exchange between different motifs and drift the system towards more stable interaction patterns and ordering. Importantly, we also show that increasing linker dynamics can compensate the effect of rigidifying mutations and prohibit aggregation. Development of a higher-resolution model, based on the same mathematical framework is undergoing in our laboratory.

## Acknowledgement

Financial support is provided by GINOP-2.3.2-15-2016-00044, HAS 11015 and the DE Excellence Program (M.F.).

---

**Figure S1 Fibrillization as a function of M3 motif affinity.** The propensity of polymers with amyloid-like elements is shown as a function of M3 motif affinity during step= 10000 – 15000 of the trajectory. Affinity of all other motifs was set to S=0.50.

## References

[1] H. Wu, M. Fuxreiter, The Structure and Dynamics of Higher-Order Assemblies: Amyloids, Signalosomes, and Granules, Cell, 165 (2016) 1055–1066.

[2] R. Nelson, M.R. Sawaya, M. Balbirnie, A.O. Madsen, C. Riekel, R. Grothe, D. Eisenberg, Structure of the cross-beta spine of amyloid-like fibrils, Nature, 435 (2005) 773–778.

[3] M.R. Sawaya, S. Sambashivan, R. Nelson, M.I. Ivanova, S.A. Sievers, M.I. Apostol, M.J. Thompson, M. Balbirnie, J.J. Wiltzius, H.T. McFarlane, A.O. Madsen, C. Riekel, D. Eisenberg, Atomic structures of amyloid cross-beta spines reveal varied steric zippers, Nature, 447 (2007) 453–457.

[4] E. Boke, M. Ruer, M. Wuhr, M. Coughlin, R. Lemaitre, S.P. Gygi, S. Alberti, D. Drechsel, A.A. Hyman, T.J. Mitchison, Amyloid-like Self-Assembly of a Cellular Compartment, Cell, 166 (2016) 637–650.

[5] C.P. Brangwynne, C.R. Eckmann, D.S. Courson, A. Rybarska, C. Hoege, J. Gharakhani, F. Julicher, A.A. Hyman, Germline P granules are liquid droplets that localize by controlled dissolution/condensation, Science, 324 (2009) 1729–1732.

[6] S.F. Banani, H.O. Lee, A.A. Hyman, M.K. Rosen, Biomolecular condensates: organizers of cellular biochemistry, Nat Rev Mol Cell Biol, 18 (2017) 285–298.

[7] K.A. Burke, A.M. Janke, C.L. Rhine, N.L. Fawzi, Residue-by-Residue View of In Vitro FUS Granules that Bind the C-Terminal Domain of RNA Polymerase II, Mol Cell, 60 (2015) 231–241.

[8] R.M. Vernon, P.A. Chong, B. Tsang, T.H. Kim, A. Bah, P. Farber, H. Lin, J.D. Forman-Kay, Pi-Pi contacts are an overlooked protein feature relevant to phase separation, eLife, 7 (2018).

[9] M. Bienz, Signalosome assembly by domains undergoing dynamic head-to-tail polymerization, Trends Biochem Sci, 39 (2014) 487–495.

[10] H. Wu, Higher-order assemblies in a new paradigm of signal transduction, Cell, 153 (2013) 287–292.

[11] L. Balagopalan, R.L. Kortum, N.P. Coussens, V.A. Barr, L.E. Samelson, The Linker for Activation of T Cells (LAT) Signaling Hub: From Signaling Complexes to Microclusters, J Biol Chem, 290 (2015) 26422–26429.

[12] J.A. West, M. Mito, S. Kurosaka, T. Takumi, C. Tanegashima, T. Chujo, K. Yanaka, R.E. Kingston, T. Hirose, C. Bond, A. Fox, S. Nakagawa, Structural, super-resolution microscopy analysis of paraspeckle nuclear body organization, J Cell Biol, 214 (2016) 817–830.

[13] D.M. Mitrea, J.A. Cika, C.B. Stanley, A. Nourse, P.L. Onuchic, P.R. Banerjee, A.H. Phillips, C.G. Park, A.A. Deniz, R.W. Kriwacki, Self-interaction of NPM1 modulates multiple mechanisms of liquid-liquid phase separation, Nature communications, 9 (2018) 842.

[14] S.F. Mitchell, R. Parker, Principles and properties of eukaryotic mRNPs, Mol Cell, 54 (2014) 547–558.

[15] P. Anderson, N. Kedersha, P. Ivanov, Stress granules, P-bodies and cancer, Biochim Biophys Acta, 1849 (2015) 861–870.

[16] T.J. Nott, E. Petsalaki, P. Farber, D. Jervis, E. Fussner, A. Plochowietz, T.D. Craggs, D.P. Bazett-Jones, T. Pawson, J.D. Forman-Kay, A.J. Baldwin, Phase transition of a disordered nuage protein generates environmentally responsive membraneless organelles, Mol Cell, 57 (2015) 936–947.

[17] S. Boeynaems, S. Alberti, N.L. Fawzi, T. Mittag, M. Polymenidou, F. Rousseau, J. Schymkowitz, J. Shorter, B. Wolozin, L. Van Den Bosch, P. Tompa, M. Fuxreiter, Protein Phase Separation: A New Phase in Cell Biology, Trends Cell Biol, 28 (2018) 420–435.

[18] N. Kedersha, M.R. Cho, W. Li, P.W. Yacono, S. Chen, N. Gilks, D.E. Golan, P. Anderson, Dynamic shuttling of TIA-1 accompanies the recruitment of mRNA to mammalian stress granules, J Cell Biol, 151 (2000) 1257–1268.

[19] S. Milles, D. Mercadante, I.V. Aramburu, M.R. Jensen, N. Banterle, C. Koehler, S. Tyagi, J. Clarke, S.L. Shammas, M. Blackledge, F. Grater, E.A. Lemke, Plasticity of an Ultrafast Interaction between Nucleoporins and Nuclear Transport Receptors, Cell, 163 (2015) 734–745.

[20] J.S. Andersen, Y.W. Lam, A.K. Leung, S.E. Ong, C.E. Lyon, A.I. Lamond, M. Mann, Nucleolar proteome dynamics, Nature, 433 (2005) 77–83.

[21] F. Wippich, B. Bodenmiller, M.G. Trajkovska, S. Wanka, R. Aebersold, L. Pelkmans, Dual specificity kinase DYRK3 couples stress granule condensation/dissolution to mTORC1 signaling, Cell, 152 (2013) 791–805.

[22] E. Gomes, J. Shorter, The molecular language of membraneless organelles, J Biol Chem, (2018).

[23] S. Jain, J.R. Wheeler, R.W. Walters, A. Agrawal, A. Barsic, R. Parker, ATPase-Modulated Stress Granules Contain a Diverse Proteome and Substructure, Cell, 164 (2016) 487–498.

[24] A.A. Hyman, C.A. Weber, F. Julicher, Liquid-liquid phase separation in biology, Annu Rev Cell Dev Biol, 30 (2014) 39–58.

[25] P.J. Flory, Thermodynamics of high polymer solutions, The Journal of chemical physics, 10 (1942) 51–61.

[26] C.P. Brangwynne, P. Tompa, R.V. Pappu, Polymer physics of intracellular phase transitions, Nature Physics, 11 (2015) 899–904.

[27] J. Wang, J.M. Choi, A.S. Holehouse, H.O. Lee, X. Zhang, M. Jahnel, S. Maharana, R. Lemaitre, A. Pozniakovsky, D. Drechsel, I. Poser, R.V. Pappu, S. Alberti, A.A. Hyman, A Molecular Grammar Governing the Driving Forces for Phase Separation of Prion-like RNA Binding Proteins, Cell, 174 (2018) 688–699 e616.

[28] A.N. Semenov, M. Rubinstein, Thermoreversible Gelation in Solutions of Associative Polymers. 1. Statics, Macromolecules 31 (1998) 1373–1385.

[29] P. Li, S. Banjade, H.C. Cheng, S. Kim, B. Chen, L. Guo, M. Llaguno, J.V. Hollingsworth, D.S. King, S.F. Banani, P.S. Russo, Q.X. Jiang, B.T. Nixon, M.K. Rosen, Phase transitions in the assembly of multivalent signalling proteins, Nature, 483 (2012) 336–340.

[30] S.K. Burley, G.A. Petsko, Weakly polar interactions in proteins, Adv Protein Chem, 39 (1988) 125–189.

[31] J.P. Gallivan, D.A. Dougherty, Cation-pi interactions in structural biology, Proc Natl A cad Sci U S A, 96 (1999) 9459–9464.

[32] J.P. Gallivan, D.A. Dougherty, A computational study of cation-p interaction vs salt bridges in aqueous media: implications for protein engineering., J. Am. Chem. Soc., 122 (2000) 870–874.

[33] M. Levitt, M.F. Perutz, Aromatic rings as hydrogen bond acceptors, J Mol Biol, 201 (1988) 751–754.

[34] T.S. Harmon, A.S. Holehouse, M.K. Rosen, R.V. Pappu, Intrinsically disordered linkers determine the interplay between phase separation and gelation in multivalent proteins, eLife, 6 (2017).

[35] P.R. ten Wolde, D. Frenkel, Enhancement of protein crystal nucleation by critical density fluctuations, Science, 277 (1997) 1975–1978.

[36] A. Molliex, J. Temirov, J. Lee, M. Coughlin, A.P. Kanagaraj, H.J. Kim, T. Mittag, J.P. Taylor, Phase Separation by Low Complexity Domains Promotes Stress Granule Assembly and Drives Pathological Fibrillization, Cell, 163 (2015) 123–133.

[37] A. Patel, H.O. Lee, L. Jawerth, S. Maharana, M. Jahnel, M.Y. Hein, S. Stoynov, J. Mahamid, S. Saha, T.M. Franzmann, A. Pozniakovski, I. Poser, N. Maghelli, L.A. Royer, M. Weigert, E.W. Myers, S. Grill, D. Drechsel, A.A. Hyman, S. Alberti, A Liquid-to-Solid Phase Transition of the ALS Protein FUS Accelerated by Disease Mutation, Cell, 162 (2015) 1066–1077.

[38] T.P. Knowles, M. Vendruscolo, C.M. Dobson, The amyloid state and its association with protein misfolding diseases, Nat Rev Mol Cell Biol, 15 (2014) 384–396.

[39] M. Ramaswami, J.P. Taylor, R. Parker, Altered ribostasis: RNA-protein granules in degenerative disorders, Cell, 154 (2013) 727–736.

[40] H.J. Kim, N.C. Kim, Y.D. Wang, E.A. Scarborough, J. Moore, Z. Diaz, K.S. MacLea, B. Freibaum, S. Li, A. Molliex, A.P. Kanagaraj, R. Carter, K.B. Boylan, A.M. Wojtas, R. Rademakers, J.L. Pinkus, S.A. Greenberg, J.Q. Trojanowski, B.J. Traynor, B.N. Smith, S. Topp, A.S. Gkazi, J. Miller, C.E. Shaw, M. Kottlors, J. Kirschner, A. Pestronk, Y.R. Li, A.F. Ford, A.D. Gitler, M. Benatar, O.D. King, V.E. Kimonis, E.D. Ross, C.C. Weihl, J. Shorter, J.P. Taylor, Mutations in prion-like domains in hnRNPA2B1 and hnRNPA1 cause multisystem proteinopathy and ALS, Nature, 495 (2013) 467–473.

[41] I.R. Mackenzie, A.M. Nicholson, M. Sarkar, J. Messing, M.D. Purice, C. Pottier, K. Annu, M. Baker, R.B. Perkerson, A. Kurti, B.J. Matchett, T. Mittag, J. Temirov, G.R. Hsiung, C. Krieger, M.E. Murray, M. Kato, J.D. Fryer, L. Petrucelli, L. Zinman, S. Weintraub, M. Mesulam, J. Keith, S.A. Zivkovic, V. Hirsch-Reinshagen, R.P. Roos, S. Zuchner, N.R. Graff-Radford, R.C. Petersen, R.J. Caselli, Z.K. Wszolek, E. Finger, C. Lippa, D. Lacomis, H. Stewart, D.W. Dickson, H.J. Kim, E. Rogaeva, E. Bigio, K.B. Boylan, J.P. Taylor, R. Rademakers, TIA1 Mutations in Amyotrophic Lateral Sclerosis and Frontotemporal Dementia Promote Phase Separation and Alter Stress Granule Dynamics, Neuron, 95 (2017) 808–816 e809.

[42] J. Sreedharan, I.P. Blair, V.B. Tripathi, X. Hu, C. Vance, B. Rogelj, S. Ackerley, J.C. Durnall, K.L. Williams, E. Buratti, F. Baralle, J. de Belleroche, J.D. Mitchell, P.N. Leigh, A. Al-Chalabi, C.C. Miller, G. Nicholson, C.E. Shaw, TDP-43 mutations in familial and sporadic amyotrophic lateral sclerosis, Science, 319 (2008) 1668–1672.

[43] S. Ambadipudi, J. Biernat, D. Riedel, E. Mandelkow, M. Zweckstetter, Liquid-liquid phase separation of the microtubule-binding repeats of the Alzheimer-related protein Tau, Nature communications, 8 (2017) 275.

[44] B. Tuu-Szabo, L. Koczy, M. Fuxreiter, Simulations of Higher-Order Protein Organizations Using a Fuzzy Framework, Complexity, 2018 (2018) Article ID 6360846.

[45] F. Troilo, C. Bignon, S. Gianni, M. Fuxreiter, S. Longhi, Experimental Characterization of Fuzzy Protein Assemblies: Interactions of Paramyxoviral NTAIL Domains With Their Functional Partners, Methods Enzymol, 611 (2018) 137–192.

[46] J. Kragelj, V. Ozenne, M. Blackledge, M.R. Jensen, Conformational propensities of intrinsically disordered proteins from NMR chemical shifts, Chemphyschem, 14 (2013) 3034–3045.

[47] M. Fuxreiter, Fold or not to fold upon binding - does it really matter?, Current Opinion in Structural Biology, 54 (2018) 19–25.

[48] P. Sormanni, D. Piovesan, G.T. Heller, M. Bonomi, P. Kukic, C. Camilloni, M. Fuxreiter, Z. Dosztanyi, R.V. Pappu, M.M. Babu, S. Longhi, P. Tompa, A.K. Dunker, V.N. Uversky, S.C. Tosatto, M. Vendruscolo, Simultaneous quantification of protein order and disorder, Nat Chem Biol, 13 (2017) 339–342.

[49] M.E. Oates, P. Romero, T. Ishida, M. Ghalwash, M.J. Mizianty, B. Xue, Z. Dosztanyi, V.N. Uversky, Z. Obradovic, L. Kurgan, A.K. Dunker, J. Gough, D(2)P(2): database of disordered protein predictions, Nucleic Acids Res, 41 (2013) D508–516.

[50] L.A. Zadeh, Fuzzy sets, Information and Control, 8 (1965) 338–353.

[51] S.R. Ainavarapu, J. Brujic, H.H. Huang, A.P. Wiita, H. Lu, L. Li, K.A. Walther, M. Carrion-Vazquez, H. Li, J.M. Fernandez, Contour length and refolding rate of a small protein controlled by engineered disulfide bonds, Biophys J, 92 (2007) 225–233.

[52] D. Brune, S. Kim, Predicting protein diffusion coefficients, Proc Natl A cad Sci U S A, 90 (1993) 3835–3839.

[53] A. De Luca, S. Termini, A definition of nonprobabilistic entropy in the setting of fuzzy set theory., Inf. Control, 20 (1972) 301–312.

[54] M. Fuxreiter, Fuzziness in Protein Interactions-A Historical Perspective, J Mol Biol, 430 (2018) 2278–2287.

[55] M. Fuxreiter, Towards a Stochastic Paradigm: From Fuzzy Ensembles to Cellular Functions, Molecules, 23 (2018).

[56] Y. Lin, D.S. Protter, M.K. Rosen, R. Parker, Formation and Maturation of Phase-Separated Liquid Droplets by RNA-Binding Proteins, Mol Cell, 60 (2015) 208–219.

[57] S. Banjade, Q. Wu, A. Mittal, W.B. Peeples, R.V. Pappu, M.K. Rosen, Conserved interdomain linker promotes phase separation of the multivalent adaptor protein Nck, Proc Natl A cad Sci U S A, 112 (2015) E6426–6435.

[58] D.T. Murray, M. Kato, Y. Lin, K.R. Thurber, I. Hung, S.L. McKnight, R. Tycko, Structure of FUS Protein Fibrils and Its Relevance to Self-Assembly and Phase Separation of Low-Complexity Domains, Cell, 171 (2017) 615–627 e616.

[59] T. Murakami, S. Qamar, J.Q. Lin, G.S. Schierle, E. Rees, A. Miyashita, A.R. Costa, R.B. Dodd, F.T. Chan, C.H. Michel, D. Kronenberg-Versteeg, Y. Li, S.P. Yang, Y. Wakutani, W. Meadows, R.R. Ferry, L. Dong, G.G. Tartaglia, G. Favrin, W.L. Lin, D.W. Dickson, M. Zhen, D. Ron, G. Schmitt-Ulms, P.E. Fraser, N.A. Shneider, C. Holt, M. Vendruscolo, C.F. Kaminski, P. St George-Hyslop, ALS/FTD Mutation-Induced Phase Transition of FUS Liquid Droplets and Reversible Hydrogels into Irreversible Hydrogels Impairs RNP Granule Function, Neuron, 88 (2015) 678–690.

[60] C. Ader, S. Frey, W. Maas, H.B. Schmidt, D. Gorlich, M. Baldus, Amyloid-like interactions within nucleoporin FG hydrogels, Proc Natl Acad Sci U S A, 107 (2010) 6281–6285.

[61] R. Halfmann, J.R. Wright, S. Alberti, S. Lindquist, M. Rexach, Prion formation by a yeast GLFG nucleoporin, Prion, 6 (2012) 391–399.

[62] S. Xiang, M. Kato, L.C. Wu, Y. Lin, M. Ding, Y. Zhang, Y. Yu, S.L. McKnight, The LC Domain of hnRNPA2 Adopts Similar Conformations in Hydrogel Polymers, Liquid-like Droplets, and Nuclei, Cell, 163 (2015) 829–839.

[63] L. Malinovska, S. Kroschwald, S. Alberti, Protein disorder, prion propensities, and self-organizing macromolecular collectives, Biochim Biophys Acta, 1834 (2013) 918–931.

[64] M.P. Hughes, M.R. Sawaya, D.R. Boyer, L. Goldschmidt, J.A. Rodriguez, D. Cascio, L. Chong, T. Gonen, D.S. Eisenberg, Atomic structures of low-complexity protein segments reveal kinked beta sheets that assemble networks, Science, 359 (2018) 698–701.

[65] H.B. Schmidt, D. Gorlich, Nup98 FG domains from diverse species spontaneously phase-separate into particles with nuclear pore-like permselectivity, eLife, 4 (2015).

[66] P.R. Banerjee, A.N. Milin, M.M. Moosa, P.L. Onuchic, A.A. Deniz, Reentrant Phase Transition Drives Dynamic Substructure Formation in Ribonucleoprotein Droplets, Angew Chem Int Ed Engl, 56 (2017) 11354–11359.

[67] Z. Monahan, V.H. Ryan, A.M. Janke, K.A. Burke, S.N. Rhoads, G.H. Zerze, R. O’Meally, G.L. Dignon, A.E. Conicella, W. Zheng, R.B. Best, R.N. Cole, J. Mittal, F. Shewmaker, N.L. Fawzi, Phosphorylation of the FUS low-complexity domain disrupts phase separation, aggregation, and toxicity, EMBO J, 36 (2017) 2951–2967.

[68] E. Delaforge, J. Kragelj, L. Tengo, A. Palencia, S. Milles, G. Bouvignies, N. Salvi, M. Blackledge, M.R. Jensen, Deciphering the Dynamic Interaction Profile of an Intrinsically Disordered Protein by NMR Exchange Spectroscopy, J Am Chem Soc, 140 (2018) 1148–1158.

[69] P.H. Nguyen, M.S. Li, G. Stock, J.E. Straub, D. Thirumalai, Monomer adds to preformed structured oligomers of Abeta-peptides by a two-stage dock-lock mechanism, Proc Natl A cad Sci U S A, 104 (2007) 111–116.

[70] J. Dogan, S. Gianni, P. Jemth, The binding mechanisms of intrinsically disordered proteins, Phys Chem Chem Phys, 16 (2014) 6323–6331.

[71] S. Hadzi, A. Mernik, C. Podlipnik, R. Loris, J. Lah, The Thermodynamic Basis of the Fuzzy Interaction of an Intrinsically Disordered Protein, Angew Chem Int Ed Engl, 56 (2017) 14494–14497.

[72] L.M. Tuttle, D. Pacheco, L. Warfield, J. Luo, J. Ranish, S. Hahn, R.E. Klevit, Gcn4-Mediator Specificity Is Mediated by a Large and Dynamic Fuzzy Protein-Protein Complex, Cell reports, 22 (2018) 3251–3264.

[73] M. Dosnon, D. Bonetti, A. Morrone, J. Erales, E. di Silvio, S. Longhi, S. Gianni, Demonstration of a Folding after Binding Mechanism in the Recognition between the Measles Virus NTAIL and X Domains, ACS chemical biology, 10 (2015) 795–802.

[74] D. Bonetti, F. Troilo, M. Brunori, S. Longhi, S. Gianni, How Robust Is the Mechanism of Folding-Upon-Binding for an Intrinsically Disordered Protein?, Biophys J, 114 (2018) 1889–1894.

[75] A. Toto, C. Camilloni, R. Giri, M. Brunori, M. Vendruscolo, S. Gianni, Molecular Recognition by Templated Folding of an Intrinsically Disordered Protein, Scientific reports, 6 (2016) 21994.

[76] M. Miskei, C. Antal, M. Fuxreiter, FuzDB: database of fuzzy complexes, a tool to develop stochastic structure-function relationships for protein complexes and higher-order assemblies, Nucleic Acids Res, 45 (2017) D228–D235.

[77] M. Arbesu, G. Iruela, H. Fuentes, J.M.C. Teixeira, M. Pons, Intramolecular Fuzzy Interactions Involving Intrinsically Disordered Domains, Front Mol Biosci, 5 (2018) 39.

[78] Y. Xiao, B. Ma, D. McElheny, S. Parthasarathy, F. Long, M. Hoshi, R. Nussinov, Y. Ishii, Abeta(1-42) fibril structure illuminates self-recognition and replication of amyloid in Alzheimer’s disease, Nat Struct Mol Biol, 22 (2015) 499–505.

[79] F. Hou, L. Sun, H. Zheng, B. Skaug, Q.X. Jiang, Z.J. Chen, MAVS forms functional prion-like aggregates to activate and propagate antiviral innate immune response, Cell, 146 (2011) 448–461.

[80] R. Ciuffa, T. Lamark, A.K. Tarafder, A. Guesdon, S. Rybina, W.J. Hagen, T. Johansen, C. Sachse, The selective autophagy receptor p62 forms a flexible filamentous helical scaffold, Cell reports, 11 (2015) 748–758.

[81] S. Frey, R.P. Richter, D. Gorlich, FG-rich repeats of nuclear pore proteins form a three-dimensional meshwork with hydrogel-like properties, Science, 314 (2006) 815–817.

