## Supplementary figures and images for "Altered dynamics may drift pathological fibrillization in membraneless organelles"

### Supplemental Figure 1

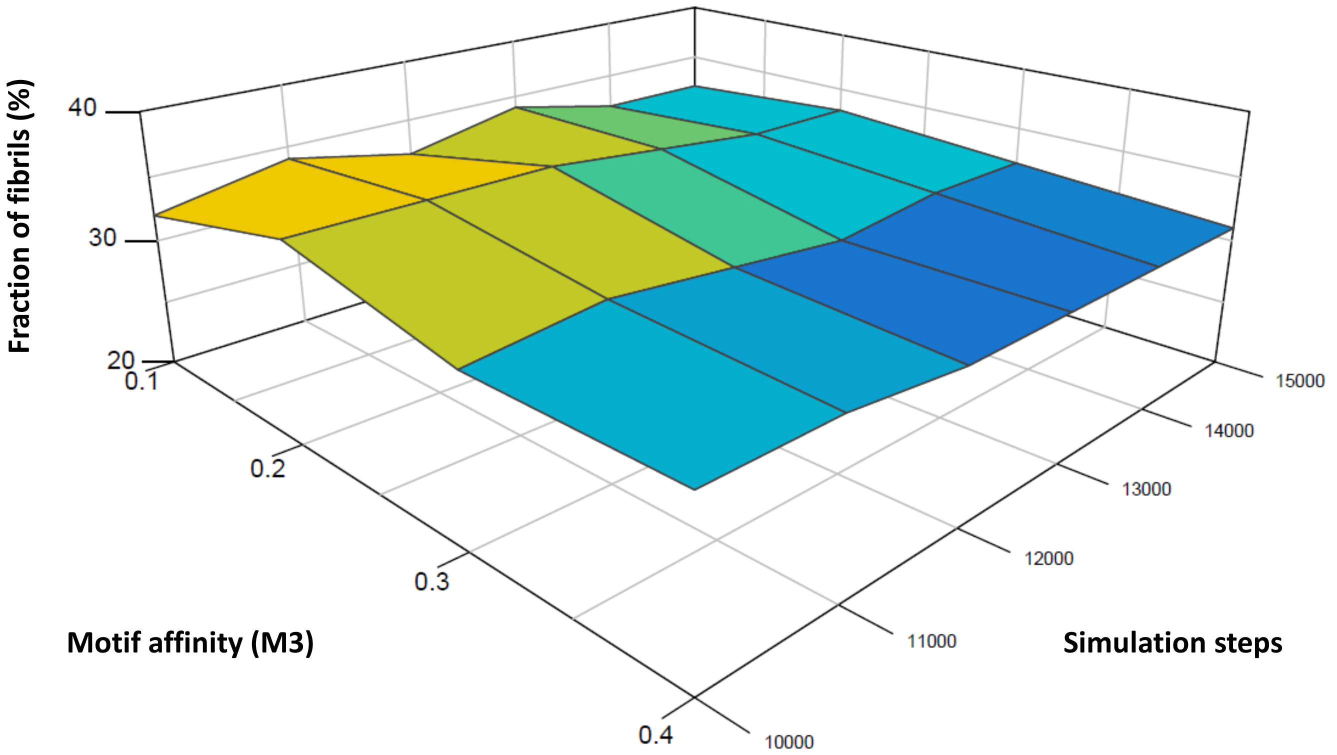
